# Mammal niches are not conserved over continental scales

**DOI:** 10.1101/2025.01.17.633640

**Authors:** Benjamin R. Goldstein, Krishna Pacifici, Alex J. Jensen, Brian J. Reich, Michael V. Cove, William J. McShea, Brigit Rooney, Elizabeth M. Kierepka, Rachel A. Roberts, Roland Kays

**Affiliations:** Department of Forestry and Environmental Resources, North Carolina State University. 2720 Faucette Dr, Raleigh, NC, USA; North Carolina Museum of Natural Sciences. 11 W Jones St, Raleigh, NC, USA; Department of Statistics, North Carolina State University; Smithsonian’s National Zoo and Conservation Biology Institute. 1500 Remount Rd, Front Royal, VA, USA; California Department of Fish and Wildlife. 1010 Riverside Parkway, West Sacramento, CA, USA

## Abstract

The niche conservatism hypothesis states that a species’ relationships to habitat and climate conditions are maintained across space and time^1–6^. Niche stationarity is assumed when ecologists estimate species’ habitat needs or transfer findings across geographic regions^7,8^. Recent studies show that some species’ associations with climate and habitat vary spatially, contradicting the niche conservatism hypothesis^9–18^. The sources of this nonstationarity are unknown and potential mechanisms remain untested. Here we show that the environmental niches of 36 common North American mammals vary spatially across dimensions of human influence, climate, and landscape. Spatial variation is not explained by known genetic subspecies lineages. Instead, niches vary at a relatively fine spatial scale consistent with adaptations by animals to local conditions, which may be explained by genetic or behavioral changes or by unmodeled interactions with unobserved variables. Spatially varying ecology means that static niche models are not appropriate at large scales and extrapolating species’ niches to novel contexts may be impossible^19^. This complicates management decisions based on transferring findings across space and projecting species’ current environmental associations to responses to future change, but may provide optimism if nonstationarity represents potential for rapid evolutionary rescue.

## Main

The habitat, climate, and geographical conditions to which a species is adapted define its “niche”^1,2^. An extension of the niche concept, known as “phylogenetic niche conservatism,” holds that a species’ niche should be maintained across space and time^3–6,20^. This assumption that the niche is spatially stationary is commonly applied in ecological practice such as when modeling species’ distributions by assuming that a species’ environmental relationships are the same across its range^7,21,22^. In conservation management, the assumption that niches are stable in space is also used to forecast species’ distributions under novel conditions, such as under climate change or for invasive species^8,23–25^, and to extrapolate knowledge of a species’ ecology from one population to inform the management of another population of the same species^26,27^. However, evidence has emerged suggesting that some niches may be nonstationary^9–12,15,16^. Higher resolution species occurrence data and advances in statistical methods for quantifying spatially varying effects have buoyed new efforts to investigate niche conservatism^11,13,15,28–31^. Models representing spatial nonstationarity in the niche with spatially varying coefficients (SVCs) have been found to outperform stationary models according to measures of predictive accuracy or parsimony^10,15^. Spatially varying species-environment relationships are therefore biologically plausible and evident in some recent models, challenging the assumptions that ecologists depend on to quantify the niche and extrapolate species distributions. As human intervention triggers rapid changes in habitat and climate, exposing many species to novel combinations of environmental conditions^19^, ecologists urgently need to resolve this issue.

Two classes of phenomena could explain apparent nonstationarity in the niche: (1) genetic or behavioral adaptation, i.e., differences in the animals themselves^32,33^; or (2) context dependence, i.e., differences in the meaning of environmental variables across the species’ range^10,34,35^. In the first case, biological differences between animals in different populations across the range lead to distinct environmental requirements for the species. Organismal differences could occur at large spatial scales^33,36^ due to ancient genetic differentiation (i.e., phylogenetic lineages), in which case species distribution models could be made for each genetic group^37^. Alternatively, organismal differences could occur at small spatial scales due to rapid adaptive evolution^38^, which could be captured by comparatively fine-scale SVCs. The second phenomenon, mechanistic context dependence^35^, could arise if the meaning of a niche dimension varies spatially. For example, a species might respond to “percent forest cover” differently depending on the type of forest or on the importance of forest cover for thermoregulation in a particular climate^14^. Without an understanding of the functional form of context dependence, stationary models would be less accurate and predictions of how the species might respond to a novel environment would be incorrect.

Here, we conducted a test of the biological mechanisms underlying nonstationarity in species niches. Using state-of-the-art data integration^39,40^ and spatially explicit models, we jointly model species distributions for 36 species of North American mammals from 19,409 camera trap deployments and 1,249,912 citizen science iNaturalist observations (Figure S1). We tested for three forms of spatial structure in species-environment relationships: Pleistocene-era genetic differentiation (H1), ecoregion-level mechanistic context dependence (H2), and fine-scale local adaptation (H3) (Figure 1). North American mammals in particular offer a good opportunity to test the hypothesis that nonstationarity is a function of within-species genetic differentiation because many of these species have well established, geographically distinct subspecies genetic lineages stemming from Pleistocene refugia^36^ that span multiple physiographic regions.

**Figure 1.**
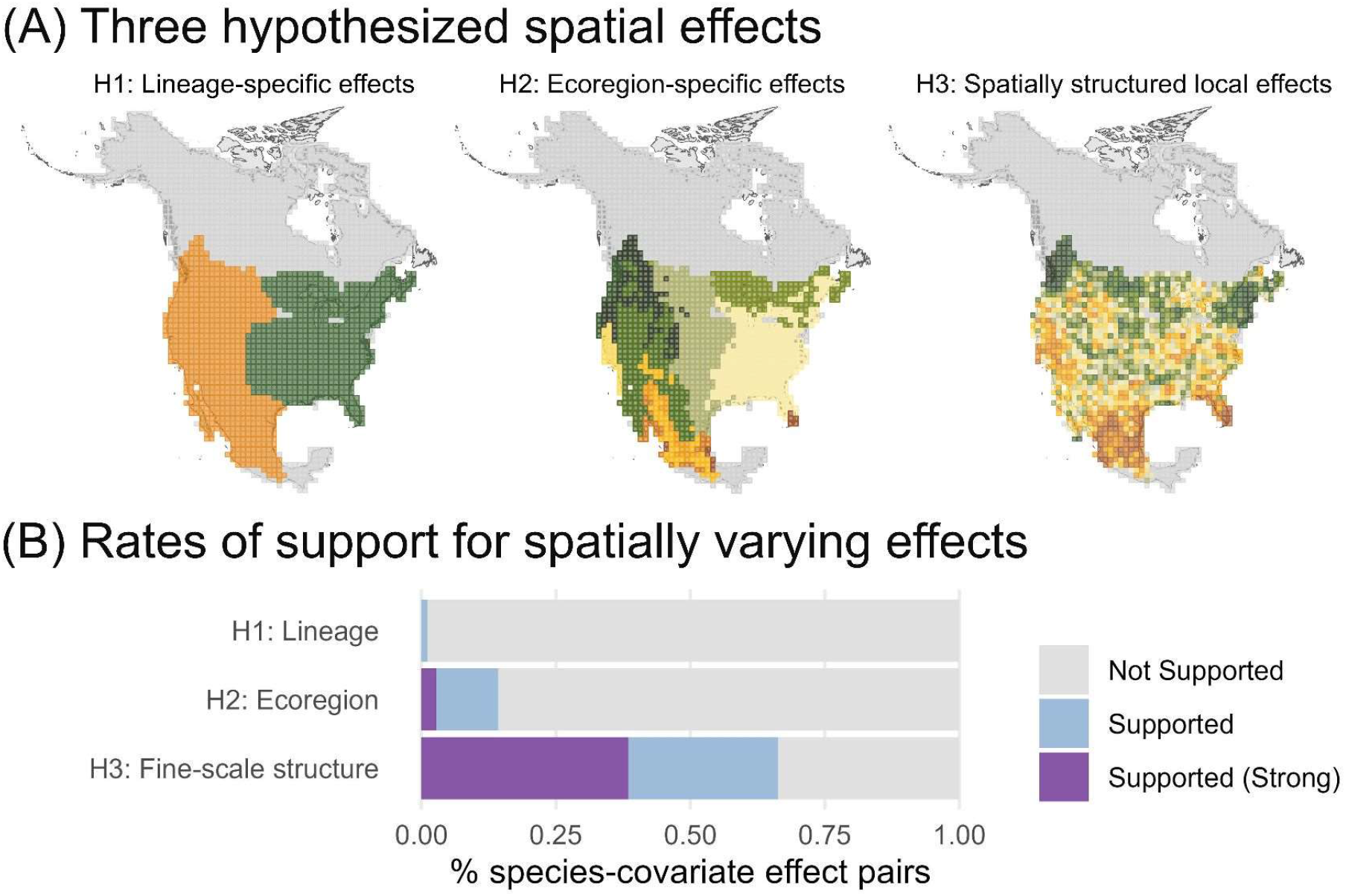
(A) Using bobcat as an example species, we display hypothetical effects within its North American range, structured according to the phylogeographic delineation of the bobcat’s two main lineages^36,41^ (H1); the 12 ecoregions it occurs in (H2); or smooth spatially structured variation (H3). (B) Rates of support for lineage, ecoregion, and fine-scale spatially varying coefficients. “Supported” indicates ≥50% posterior inclusion probability and “Supported (Strong)” indicates ≥90% posterior inclusion probability.

### Niche nonstationarity

We developed integrated species distribution models to characterize support for three types of spatially varying coefficients (SVCs) in the environmental relationships of North American mammals corresponding to three core hypotheses (Figure 1A). We considered 2 covariates of human influence (human population density and percent agriculture), 2 for climate (maximum temperature and precipitation), and 3 for landscape (terrain roughness, greenness, and percent grassland). These covariates comprise 7 dimensions of the niche considered influential for driving the spatial distribution of mammals.

Across 36 species of North American mammals, we found support for fine-scale, spatially structured SVC effects (H3) among 66.3% of species-niche dimension pairs and ecoregion effects among 14.3% (H2; Figure 1B). We found little support for larger scale genetic differences affecting ecology – among the 13 species with known Pleistocene-era subspecies genetic lineages, we found support at a rate of only 1.1% (H1; Figure 1B). SVCs were widespread across taxa, with at least one identified in all 36 species studied (mean = 5.67 supported effects per species of 14 or 21 possible per species without or with lineages; see Table S1 for species list). SVC effects were supported at comparable rates across three environmental dimensions of the niche (Figure 2B). We conducted a validation exercise and found our approach has reasonable power to detect spatially varying effects when they were truly present (true positive rate of 85.1%), even when both lineage and fine scale effects were present simultaneously. Therefore, we interpret low support for lineage-structured spatial effects as inconsistent with the hypothesis that species’ phylogenetic lineages are an important cause of spatially varying niches.

**Figure 2.**
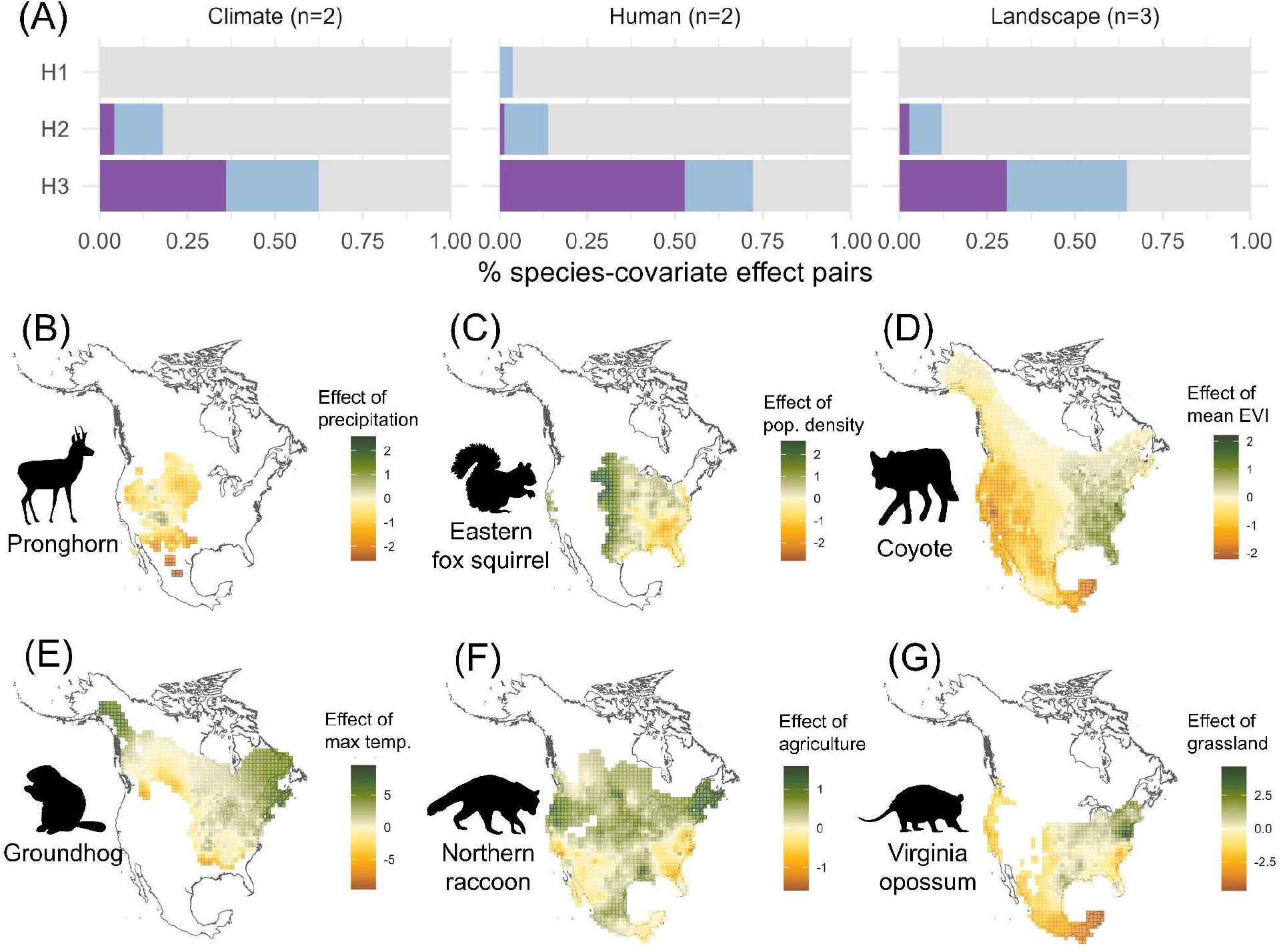
(A) The percentage of species-covariate relationships that had support (blue) or strong support (purple) for spatially varying effects summarized by covariate type. (B-G) Example maps of spatially varying covariate effects across niche dimensions. Each panel visualizes the effect of a one-standard deviation in the covariate on the relative abundance of the species within 100 km grid cells across the species’ range. For example, the abundance of Eastern fox squirrels was negatively related to human population density in the southeast, but positive along the western edge of its range (Panel C). Each example is associated with support for H3. Image silhouettes obtained on phylopic.org and are in the public domain.

### Spatial variation in the relative importance of niche dimensions

We characterized the rate at which SVC effects captured changes in the direction of the relationship between a covariate and a species’ abundance. Among species-covariate pairs with supported SVCs, we observed sign flips (i.e., effects that were positive in one part of the range and negative in another) at a rate of 16.7%, with differences in magnitude comprising the remaining 83.3% of effects (Figure S2). SVCs more commonly represent the strength of the relationship between each species and a covariate, and less commonly capture changes in the directionality of a species’ environmental relationships. The magnitude of variation was ecologically meaningful. Across all species and niche dimensions, the median difference between the 10th and 90th quantiles of the effect of a covariate on a species was 0.92 (10th quantile: 0.16, 90th quantile: 3.10), meaning that the effect of a standard deviation increase in the covariate on the species’ relative abundance was 0.92 greater in one part of the range than in another; at one standard deviation above the mean value, this translates to a 2.5x increase in relative abundance attributable to that covariate. These differences in estimated relationships lead to large downstream differences in fine-scale predictions of space use; for example, in the case of bobcat, predicted log-scale intensity varied substantially on small spatial scales, reflecting the added benefit of local estimates of habitat use (Figure 3).

**Figure 3.**
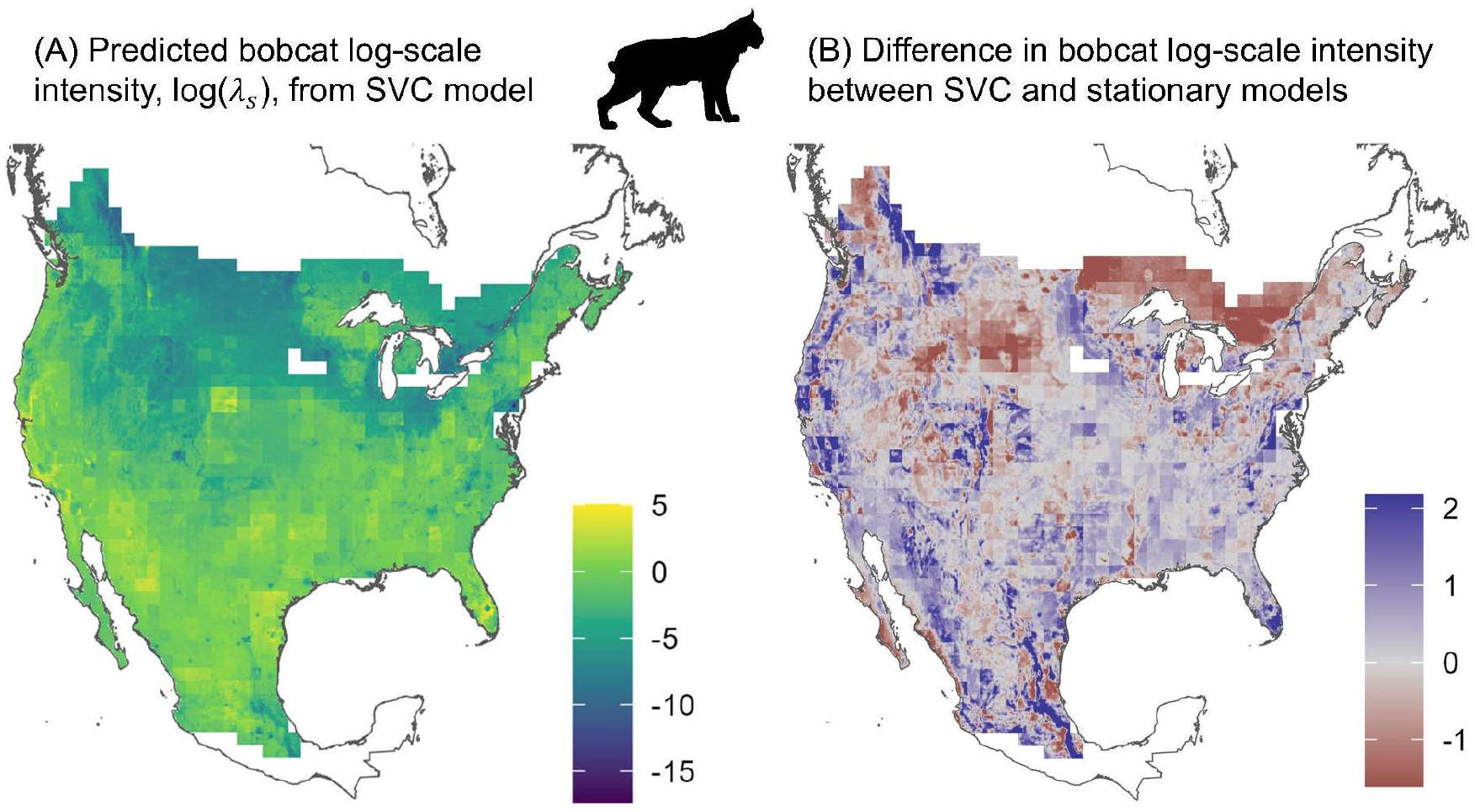
(A) Point estimates of log-scale relative abundance of bobcat across its modeled range obtained from an SVC model. (B) Comparing the predictions of intensity between the SVC model and a model that assumes stationary effects reveals large differences across the continent, illustrating the importance of accounting for the species’ spatially varying ecology when predicting fine-scale relative abundance. The no-SVC model substantially overestimated bobcat relative abundance in the north-eastern edge of its range and largely underestimated bobcat density in the south-western end of its range.

### Are SVCs capturing nonlinear covariate effects?

Because the environmental covariates modeled here are themselves spatially structured, nonlinear but spatially stationary species-environment relationships could appear in the model as spatially varying effects^35^. For example, if species consistently exhibit a quadratic relationship to temperature, a fine-scale spatially structured SVC model could estimate positive slopes in colder parts of the range and negative slopes in warmer parts of the range. This could give rise to apparent context dependence^35^. To identify whether the SVCs in our model were capturing nonlinear relationships, we investigated the extent to which the values of covariates themselves were associated with the estimated covariate effects with generalized additive models. Across 173 species-covariate pairs with supported SVCs, nonlinear relationships explained a median of 17.8% of the spatial variation in the estimated effect of that covariate (R^2^ 10th quantile = 3.8%; 90th quantile 47.6%; Figure S3). These findings suggest that SVCs may be effective at capturing nonlinear species-environment relationships that play out over large spatial scales, most commonly for climatic niches, and also capture substantial spatial variation beyond what is explained by nonlinearity.

### Model validation

We conducted a simulation study to evaluate our ability to differentiate between hypothesized SVCs, tracking the rate at which the model correctly identified support for SVCs. False positive and false negative rates were modest across all simulation conditions, meaning that our model successfully differentiated between SVC and non-SVC covariate effects in a realistic simulation (Figure S4). Using a support threshold of 50% of the posterior distribution, we observed an overall SVC false positive rate of 9.1% and a false negative rate of 14.9%. Using a strong support threshold of 90%, we observed a false positive rate of 0.7% and a false negative rate of 38.1%. Among coefficients with multiple types of SVCs operating simultaneously, we observed a false positive rate of 21.9% and a false negative rate of 17.2% at the 50% support level, and a false positive and negative rates of 1.2% and 43.0% at the 90% support level, indicating weaker but still moderate power to simultaneously identify support for competing effects. These findings suggest that our model approach has adequate power to identify spatially varying effects of reasonable magnitude.

### Implications for ecology and conservation

North American mammals exhibit different environmental associations in different parts of their range. Building on previous work finding nonstationarity at smaller scales or with single species^10–12,14^, we show that nonstationarity in the niche is found across taxonomic groups and dimensions of the niche. Our models provide new insight into the breadth of preferences exhibited by species across their ranges. For example, pronghorn may be restricted by low precipitation in the southwest (Figure 2B) while groundhog may be limited by high temperatures along the southern edges of its range (Figure 2E). Such patterns could be applied to predict where and how species are likely to be most sensitive to climate change. These models also provide meaningfully different estimates of species abundance compared to models that assume stationary niches.

We found that spatial variation in the niche was best explained by a relatively fine scale spatial process (Figure 1B). Spatial patterns in environmental relationships were not consistent with known subspecies lineages as determined by phylogeographic studies^36^, strongly indicating that these ancient genetic differences did not drive species’ spatially varying niches. We found moderate support for the hypothesis that species-environmental relationships vary by ecoregion context. Estimated effects were not associated with variation in covariates themselves, meaning that they were not attributable to a failure to properly model the functional form of covariates. The biological mechanisms driving variation in the niche are likely nuanced, but a deeper understanding is critical for informing conservation management.

The key outstanding question for differentiating between the plausible drivers of variation in the niche is: does fine-scale spatial variation in the niche reflect (1) differences in the animals themselves, or (2) mechanistic context dependence in animals’ relationships to environmental features? These two hypotheses suggest contrasting predictions for species’ responses to future change. If variation in the niche is primarily driven by fine-scale genetic differentiation, this response is likely a complex interplay between phenotypic plasticity and genomic variation, making it difficult to predict without identified candidate loci and evaluations of the strength of selection^42^. On the other hand, if variation in the niche is a result of context dependence, it is difficult to anticipate how they will respond to changing conditions: context dependence could mean either that animals may be highly plastic in their responses to environmental features or that animals’ niches may depend on interactions with unobserved features. Ultimately, differentiating between these two remaining hypotheses will require further study. Studies in local adaptation have revealed rapid evolution to anthropogenic perturbation across multiple taxa, and further genomic analysis across species’ ranges according to climatic conditions can are key to examining potential mechanisms responsible for local adaptation. Furthermore, animal translocations could provide pseudo-experimental settings for clarifying whether individuals from different populations and regions react to the same environmental conditions. If context dependence is common, ecologists could consider a multi-step modeling approach in which SVCs are used to identify the covariates and regions that are spatially variable that could then be investigated with the inclusion of missing interactions or confounding variables, enabling prediction across regions. SVCs could serve to generate hypotheses about the type of covariates associated with nonstationarity, reducing the number of potential interaction effects to be considered.

While work remains to fully explain this phenomenon, the observation that niches vary spatially has immediate and serious implications for conservation practice. First, mammal distribution models pooling data from populations more than 100 kilometers apart should consider that environmental relationships likely change across the study area. Practitioners should consider explicit SVC models^15,28^ or assess evidence of spatial structure in model residuals and thoroughly explore interaction effects between covariates. Advances in integrated modeling techniques will be critical for achieving the data volume and coverage necessary to estimate these models. Second, ecologists should not extrapolate species’ niches over long distances or to novel environmental conditions. Local or regional assessments of a species’ niche will often fail to predict the species’ niche elsewhere in its range. Ecologists should be cautious when using stationary models to inform conservation at the scale of species ranges. Instead, the field should prioritize understanding the mechanisms giving rise to spatial variation in the niche and developing models that explicitly account for those mechanisms.

## Methods

### Camera trap data

We pooled camera trap sampling effort from 19,409 camera deployments aggregated from several sources (Figure S5; Table S1). We define a deployment as a continuous sampling period associated with a single camera at a fixed location^1^. All camera traps were set at ∼0.5m height, unbaited and unpaired, and were set to take multiple pictures on each trigger with no delay between triggers. A variety of camera models were used, but all had similar fast (<0.5s) trigger time. The largest camera trap data source was the Snapshot USA Project (9,065 deployments)^2–4^ which includes resampling of many arrays across the USA from 2019-2023. Camera trap data were included from other programs, including the Carolina Critters Project^5^ (4,260 deployments, years 2016-2019), the Recreation Effects on Wildlife Project^6^ (1,894 deployments, years 2012-2013), California Department of Fish and Wildlife bobcat surveys^7^ (1,368 deployments, years 2021-2023), the Urban to Wild Project^8^ (996 deployments, years 2014-2020), and several smaller-scale sampling projects^9–17^ (Table S1 in Extended Data).

We discarded deployments shorter than 20 days or longer than 300 days. Among included deployments, the 10th quantile deployment length was 21 days, the median deployment length was 33 days, and the 90th quantile deployment length was 60 days. We summarized data obtained from each deployment as detection histories using a 10-day detection window, yielding a 0 or 1 for each complete 10-day window within each deployment indicating whether the species was detected (1) or not (0) at any point during that window^18^.

We selected for analysis 37 species that were observed on at least 100 camera deployments across at least 5 independent camera arrays (Table S2 in Extended Data). One species, the southern flying squirrel (*Glaucomys volans*), was later excluded during model fitting due to poor MCMC mixing.

### iNaturalist data

iNaturalist (iNaturalist.org) is a participatory science repository that aggregates observations of biodiversity submitted by members of the public. An observation on iNaturalist achieves “research grade” quality if several criteria are met, including having a piece of evidence (typically a photograph) that may be reviewed by other iNaturalist users, and having at least 2/3 of reviewers agree on the ID.

We downloaded all observations of mammals in North America via manual query on the iNaturalist website in January 2023 (Jensen et al., in review). The original dataset contained 1.9 million observations that occurred between June 20, 1900 and December 31, 2022. We filtered iNaturalist observations by excluding non-research grade observations and observations marked as being photographs of tracks or scat.

In each Scale 2 (50 km) grid cell, we recorded the total number of mammal observations in the filtered dataset as the cell’s “effort”, *E*_e_. Then, we recorded the number of observations of the target species in each Scale 2 grid cell as the cell’s “iNaturalist count,” *N*_e_, for that species.

### Spatial covariate data

We obtained data for seven covariates hypothesized to influence species’ intensity: two climate covariates (maximum annual daily temperature, average annual precipitation), three landscape covariates (greenness, terrain roughness, and percent grassland), and three human influence covariates (human population density and percent agriculture). Climate covariates were obtained from MERRAclim^19^. Percent grassland and percent agriculture were obtained from Jung et al.^20^. Terrain roughness was obtained from Amatulli et al.^21^. Human population density was obtained from Gridded Population of the World v4 dataset^22^. Greenness, as represented by the effective vegetation index (EVI), was obtained from the MODIS product MOD13A2.061^23^. All spatial covariates were aggregated from their original resolutions to a common 5 km by 5 km grid covering North America. We originally considered percent forest as a covariate but discarded this variable due to high multicollinearity with other variables.

We obtained two additional spatial covariates at each camera location for use in modeling detection probabilities. We calculated the distance from each camera to the nearest road using the Global Road Inventory Project v4 dataset^24^. We retrieved an estimate of canopy height at each camera from Potapov et al.^25^.

We developed species phylogeography maps for 13 species by estimating the region of their range occupied by each genetic lineage. We used the breaks between lineages digitized by Jensen et al.^26^ to partition each species’ range into regions in which we were confident in assigning to a lineage and regions where the genetic lineage was unclear. We excluded regions where lineages were unclear or unstudied from the model.

### Spatial scales and spatial extent

During modeling, we reconciled multiple spatial scales by assuming that species followed an inhomogeneous Poisson point process^27–29^ within each of 3 defined spatial scales, which we refer to throughout the Methods section. Scale 1 was a 5 km x 5 km grid on which the underlying intensity process operated (indexed in model equations by *s*). Scale 2 was a 50 km x 50 km grid over which iNaturalist observations are aggregated (indexed by *c*). Scale 3 was a 100 km x 100 km grid used to define the conditional autoregressive (CAR) spatial field (indexed by *g*).

We restricted the spatial extent of the analysis on a species-by-species basis. For 22 species without lineage information, we retrieved species ranges from IUCN^30^ and modeled only those camera deployments and iNaturalist Scale 2 cells that overlapped with the species’ range. For species with lineage information, we considered only those camera deployments and iNaturalist Scale 2 cells that overlapped subsections of the species range with clear lineages^26^.

### Integrated species distribution models

To estimate species’ range-wide relationships to environmental variables, we developed single-species integrated species distribution models (iSDMs)^27–29,31^. Camera trap detections were defined as following an occupancy model. At each camera, we discretized detections into detection histories composed of 0s and 1s representing whether the target species was observed during each consecutive 10-day detection window ^18^. The *j*th observation at camera *i, y*_ij_, was distributed according to

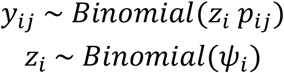

where *z*_i_ is an unobserved latent state representing whether camera *i* was used by the species during the camera’s deployment, ψ_i_ was the probability that the species used the area surveyed by the camera at all during the camera’s deployment, and *p*_ij_ was the probability of detecting the animal during the *j*th detection window. We adopted an asymptotic interpretation of occupancy, considering a given camera “occupied” if the area it surveyed was used by the species at least once during the deployment period^32^.

The camera-level occupancy probability, ψ_i_, was linked to the underlying point process model via the relationship

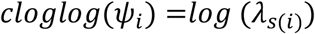

where *λ*_s(i)_ is the value of the intensity field in scale 1 grid cell *c* containing camera *i*. The detection probability associated with detection window *j* at camera *i* is defined as

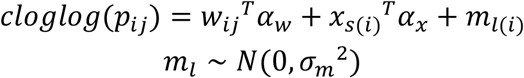

where *w*_ij_ is a vector (including an intercept) of four camera- and detection window-specific covariates with associated effects coefficients *α*_w_, *x*_s(i)_ is a vector (not including an intercept) of spatially structured covariates with effects coefficients *α*_x_, and *m*_l_ is a grouping random effect shared by detections associated with all deployments occurring at the same unique location in any year. Covariate data *x*_s(i)_ are the same as those included in the intensity submodel (see below).

The iNaturalist counts in each Scale 2 cell, *N*_e_, were modeled conditional on observed effort *E*_e_ as

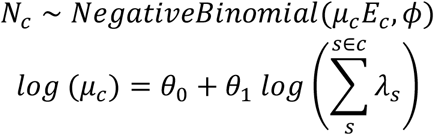

where *ϕ* is the negative binomial overdispersion parameter, *µ*_e_*E*_e_ is the mean of the negative binomial, *θ*_0_ and *θ*_1_ are parameters that control the correspondence between the intensity field and the iNaturalist data. The parameter *θ*_0_ estimates an intercept of the iNaturalist reporting rate for the species, which may be driven by human preferences or logistical obstacles to detecting the species^33,34^ but which are constant in space, while the parameter *θ*_1_ controls the strength of the correspondence between iNaturalist and the camera trap data. This model formulation treats the camera trap data as the “gold standard” dataset directly informing the latent intensity process, allowing the iNaturalist data to more weakly influence its value if the two datasets diverge^29,31^, reflecting the fact that the camera trap data are collected under a standardized protocol with known effort and less sampling bias compared to iNaturalist observations. We inspected the posterior estimates of *θ*_1_ to characterize the frequency at which the two datasets corresponded (Figure S6).

The intensity in Scale 1 cell *s, λ*_s_, is modeled as

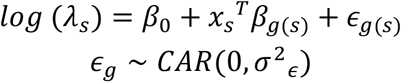

where *β*_0_ is an intercept, *x*_s_ is a vector of seven spatial covariates, *β*_g(s)_ are the spatially varying effects of the covariate, and ϵ_e(s)_ is a spatially structured random effect.

### Evaluating support for three spatially varying coefficient (SVC) mechanisms

The effect of the *i*th covariate in Scale 3 cell *c, β*_i,g_, is

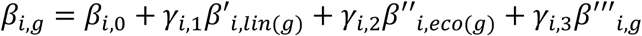

where

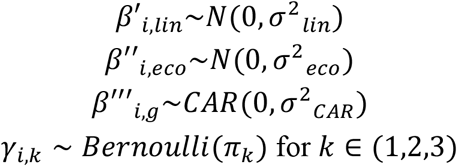

In plain language, the effect of covariate *i* on the species’ intensity in Scale 2 cell *c* is an additive combination of a spatially constant mean effect, *β*_i,0_; a random slope corresponding to the lineage associated with cell *g, β*′_i,lin(g)_; a random slope corresponding to the ecoregion in which cell *g* is found, *β*′′_i,eeo(g)_; and a zero-centered conditional autoregressive (CAR)^35^ random slope, *β*′′′_i,g_, whose distribution is determined by the adjacency matrix of the complete set of Scale 3 grid cells covering North America.

The Bernoulli switching variables γ_i,k_ turn each component of spatial variation for each covariate on (when γ_i,k_ = 1) or off (when γ_i,k_ = 0) over the spatial domain, representing selection between competing models of spatial structure in covariates^36^. If the posterior mean of γ_i,k_ was greater than a support threshold of 0.5, we characterized this as moderate support for the inclusion of the relevant spatial effect, while if it was greater than 0.9, we characterized this as strong support (see section “Simulation-based model evaluation”). For non-lineage species, we excluded the lineage random slopes from the model.

We estimated the model using Markov chain Monte Carlo (MCMC) with NIMBLE version 1.2.1^37^. To improve MCMC mixing, we curated a set of samplers, including NIMBLE’s reversible jump MCMC sampler (RJMCMC)^38^, which we applied to the indicator variables γ_i,k_. For each species, models were run for 4 chains of 20,000 iterations each including a burn-in period of 10,000 iterations. For species whose models did not converge after 4 chains, we iteratively ran 2 additional chains until convergence was achieved. Because classical convergence criteria like the Gelman-Rubin diagnostic are not appropriate for evaluating the mixing of Bernoulli random variables, which may remain at a single value (0 or 1) for long periods even in a well-mixed model, we adopted a holistic approach to evaluating model convergence. We monitored the log probability of the observed data, ***y*** and ***N***, and calculated the Gelman-Rubin diagnostic and effective sample size for this quantity to evaluate the mixing of the full model.

Aggregating across posterior iterations, we tracked the rate at which each covariate was the most influential (i.e. had the greatest estimated *β*_i,k,e_) in each cell, which we visualize in Figure 3. We then characterized each species-covariate pair with at least one supported SVC as either a “sign flip” SVC or a “non-sign flip” SVC. An SVC was denoted as a “sign flip” if it contained at least one cell with a *β*_i,e_ value with a 95% posterior credible interval (95% CI) having a lower bound greater than 0 and one having a 95% CI upper bound less than zero, indicating evidence that the covariate was positively associated with intensity in one cell and negatively associated with intensity in another. Otherwise, the SVC was denoted a “non-sign flip.” When tallying these results, we excluded species-covariate pairs without any *β*_i,k,e_ values whose posterior distributions did not overlap 0 as uninformative.

### Simulation-based model evaluation

We conducted a simulation exercise to evaluate the model’s propensity for recovering true spatial effects. In light of the large number of competing spatial processes in the model, we especially needed to evaluate the model’s ability to accurately recover multiple spatial effects. We selected two species, eastern chipmunk and bobcat, to serve as archetypes during model evaluation. We used real camera locations, iNaturalist sampling effort, and covariate values associated with each species and discarded the true detection histories and iNaturalist species counts. We simulated four datasets for each species under each of the following scenarios: (1) 4 CAR covariates, 2 lineage covariates, and 2 ecoregion covariates; (2) 2 covariates with CAR and ecoregion effects, 2 CAR covariates, and 2 lineage covariates; (3) 2 covariates with CAR and lineage effects, 2 CAR covariates, and 2 ecoregion covariates. For the full set of true parameter values used for simulation, see Table S3 in the Extended Data. We fit the main model to each of the 24 simulated datasets following the procedure outlined above. We monitored the inclusion probabilities of each covariate spatial effect and observed the false positive and false negative rates of estimated spatial effects based on posterior support thresholds of 0.5 and 0.9, i.e. whether the relevant switching variable γ_i,k_ was 1 in at least 50% or at least 90% of its posterior MCMC samples.

### SVCs and nonlinear covariate effects

We parameterized the main effects of each covariate on the intensity field as linear. While this choice is common across distribution modeling applications^39^, nonlinear species-environment relationships may sometimes occur. When covariates are spatially structured, SVCs offer an opportunity to investigate potential nonlinear relationships as they manifest across large spatial scales; for example, an SVC might estimate that a species responds more positively to temperature in colder parts of its range.

To identify whether this phenomenon was occurring in our results, we used *post hoc* analysis to characterize the extent to which variation in spatial fields was explained by nonlinear species-environment relationships. For each species-covariate pair with at least one supported SVC, we estimated a generalized additive model (GAM)^40^. Each Scale 3 spatial cell overlapping the species range was included as a point. The response value was the mean beta of the CAR field, which was regressed against a cubic regression spline with up to 50 knots on the mean value of the covariate across Scale 1 cells contained within that Scale 3 cell. We propagated error in the CAR field by weighting each point by the inverse of the variance of the effect’s posterior distribution. We interpreted the adjusted R^2^ of the GAM as a measure of the variation in the estimated covariate effect attributable to a nonlinear species-environment relationship, and tracked the distribution of R^2^-adj. values across species and covariates.

## Data availability statement

We publish a Zenodo repository^41^ with the camera trap detection data, iNaturalist observations, and covariate data at the following URL: https://doi.org/10.5281/zenodo.14219130. Data are provided as pre-processed inputs in the form needed for modeling (i.e. covariates are summarized on a 5 km grid; camera trap data are summarized to detection histories with 10-day windows; iNaturalist data are summarized to a 50 km grid; etc.). These inputs are intended for use in the fully reproducible analysis pipeline provided in the accompanying Github repository (see “Code availability statement”). This data repository also includes a file giving the estimated effect of each covariate on each species in each Scale 3 grid cell across its range, as well as raster files giving predictions of the log-scale intensity of each species, and the associated uncertainty in log-scale intensity, across the study area.

## Code availability statement

All code used to estimate the models presented can be found at the following Github repository: https://github.com/dochvam/Mammal_SVCs_ISDM_reproducible. Accompanying the code, we provide a README document outlining how to reproduce the validation simulations, model estimation procedure, and posterior visualizations presented in the paper.

## Acknowledgements

Thanks to Perry de Valpine and Sally Paganin for advising on practical challenges associated with model estimation. Thanks to the many participants in the Snapshot USA program and to the iNaturalist contributors across North America whose work and commitment to open science made this study possible. Thanks to Christopher Lopez for compiling the iNaturalist data. This work was supported by NSF Awards #2206783 and #2206784.

## Author contributions

All authors contributed to the design of the project, and all authors edited and approved of the final manuscript. KP, MVC, EMK, WJM, and RK conceived of and initiated the study and acquired funding. BRG, KP, and BJR designed the statistical methodology. BRG implemented the statistical methodology, produced visualizations, and wrote the original draft. AJJ and BR curated the data. MVC, WJM, RAR, and RK contributed original data.

