## Extended Data for "Mammal niches are not conserved over continental scales"

### Supplemental figures

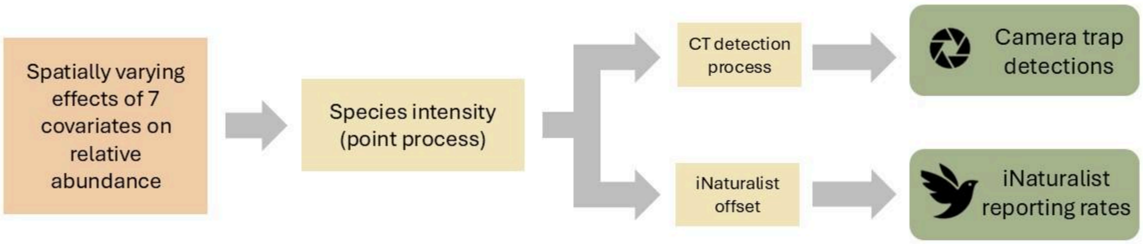

**Figure S1.** Schematic describing the integrated species distribution model designed to test three hypotheses of spatially varying species-environment relationships.

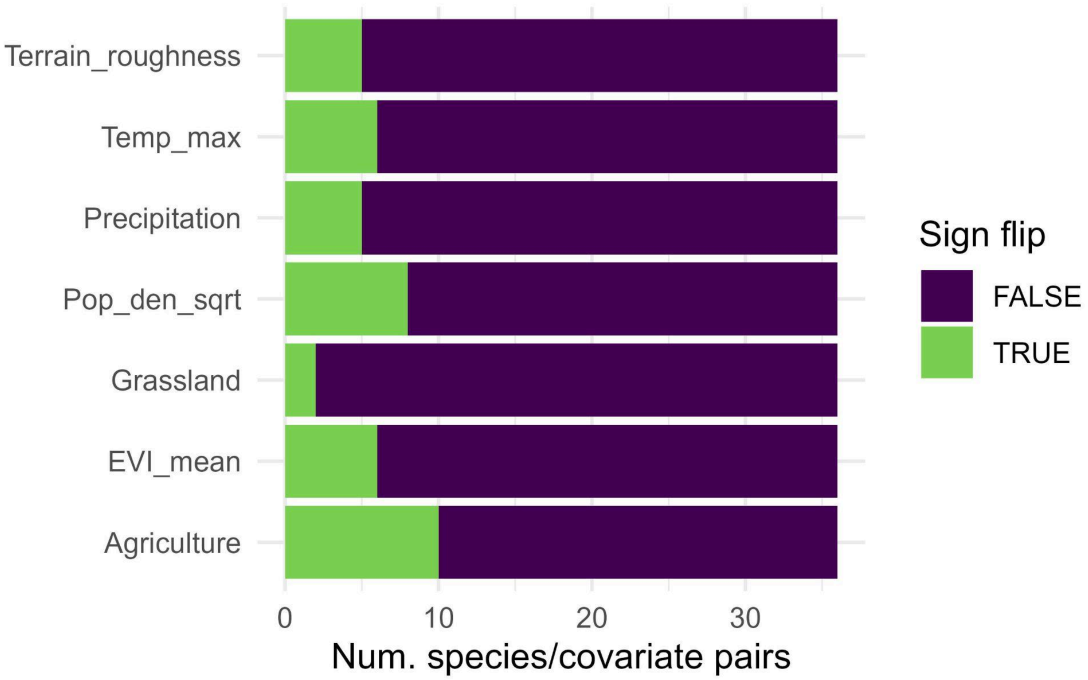

**Figure S2.** Rates of sign flips among each SVC covariate. Sign flips were relatively uncommon overall, but were most common for human population density and maximum temperature.

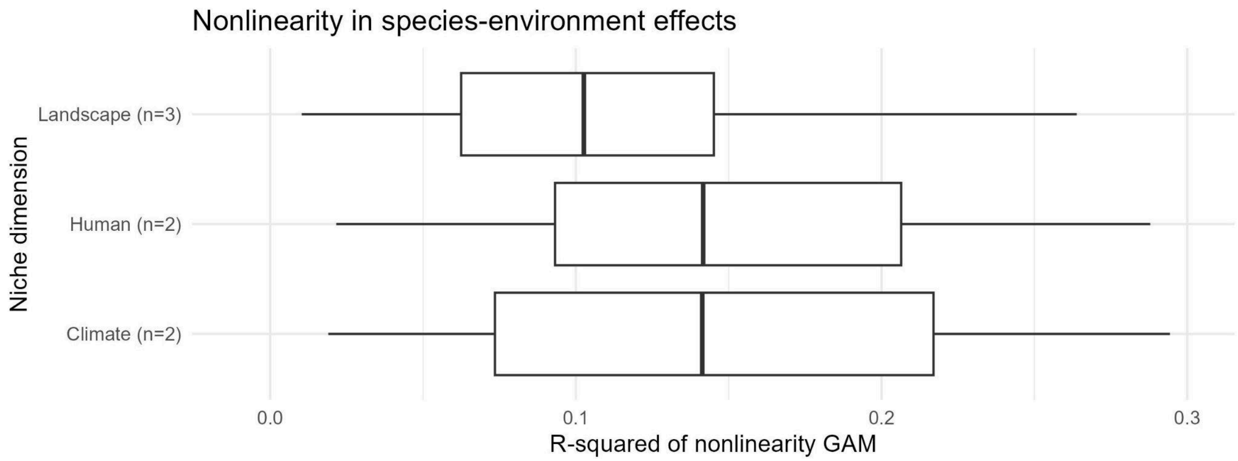

**Figure S3.** Exploring whether SVC effects represent nonlinear species-environment relationships. Across all species-covariate pairs with at least one supported SVC, variation in the value of a covariate explained only about 0-30% of spatial variation in the effect of that covariate.

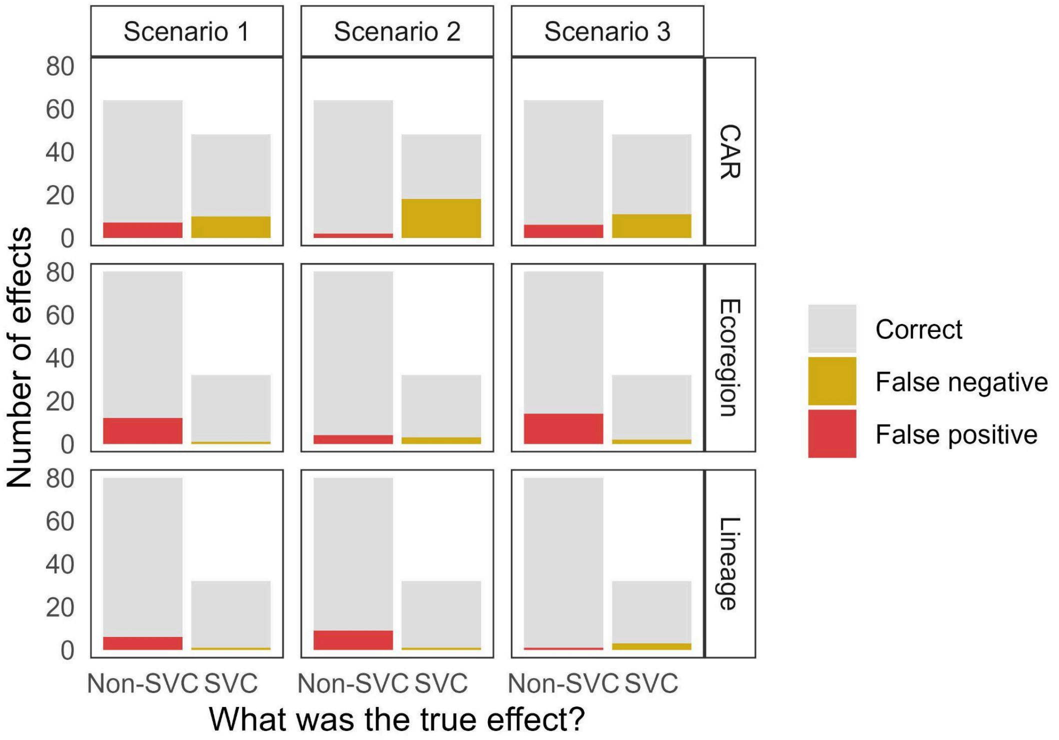

**Figure S4.** Results from a simulation-based validation exercise testing the model's ability to identify known SVC effects. This figure summarizes results across all simulated datasets based on a 50% support threshold. The overall false positive rate was 9.1% and the overall false negative rate was 14.9%. False negative rates were highest among simulated CAR effects and lowest for lineage effects.

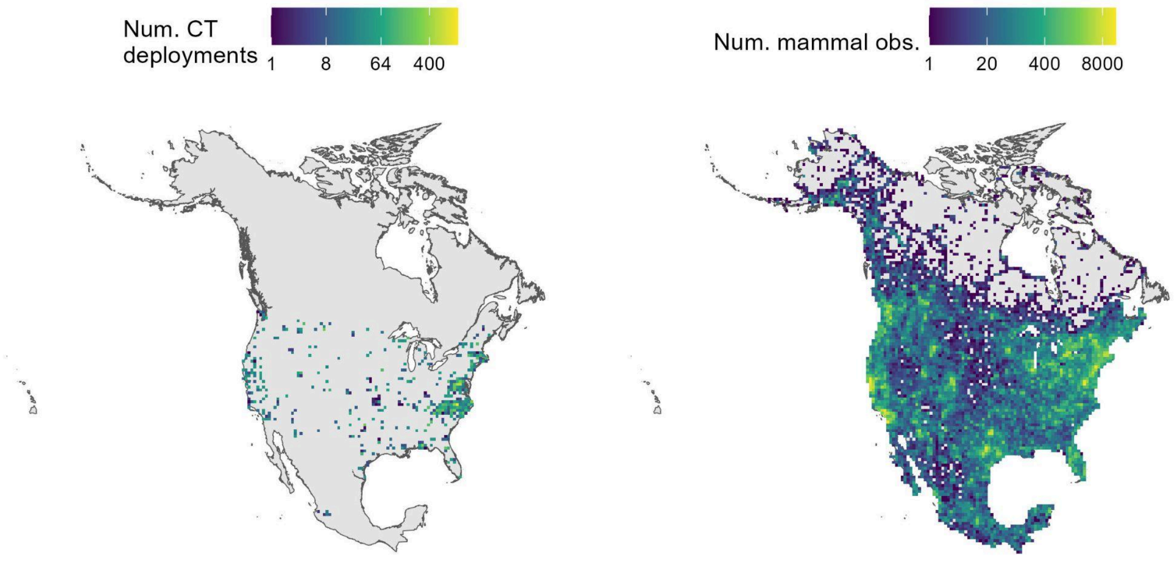

**Figure S5.** Spatial effort for 19,303 camera trap deployments (left panel) and 1,249,912 iNaturalist mammal observations (right panel) across the study area. For visualization, both datasets are aggregated to the 50 km grid used to define Scale 2.

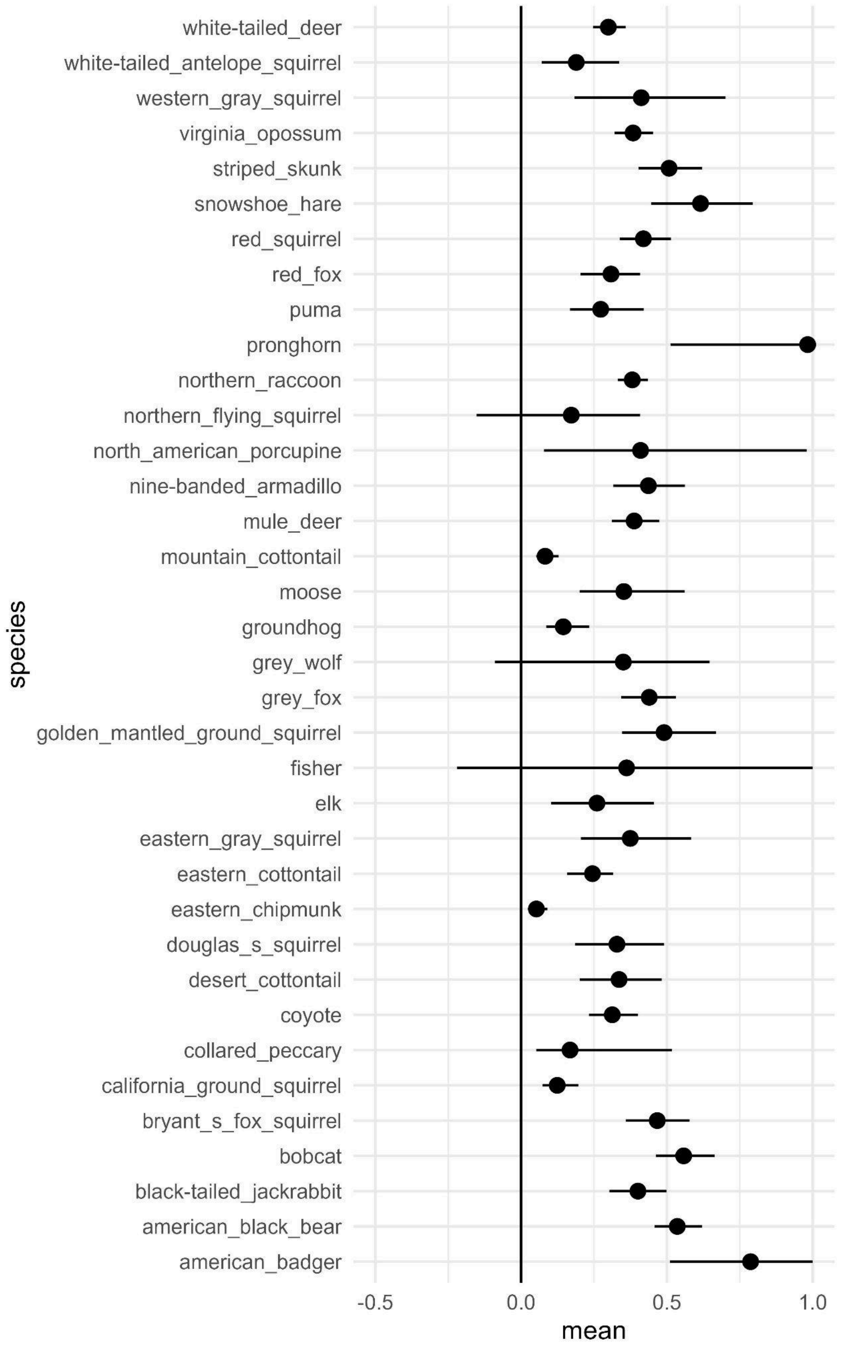

**Figure S6.** Estimated values of the join parameter $\theta_1$, which controls the relationship between iNaturalist data and the underlying intensity surface tied to camera trap occupancy. For 33 of 36 species, the 95% credible interval of the estimate of $\theta_1$ was positive and within the interval (0, 1), indicating that the iNaturalist reporting rates were positively associated with camera trap occupancy and contributed to the estimate of covariate effects.

### Table S1: Camera trap data sources

The names of each source of camera trap data included in the integrated model, along with the number of deployments associated with each source and a citation for the source. All datasets are also cited in the Methods section "Camera trap data."

| Name | Number of camera trap deployments | Citation |
|---|---|---|
| Snapshot USA | 9065 | Cove, Michael V., et al. "SNAPSHOT USA 2019: a coordinated national camera trap survey of the United States." (2021): e03353.<br><br>Kays, Roland, et al. "SNAPSHOT USA 2020: A second coordinated national camera trap survey of the United States during the COVID‐19 pandemic." (2022): e3775.<br><br>Shamon, H., et al. "SNAPSHOT USA 2021: A third coordinated national camera trap survey of the United States." Ecology, 105.6 (2024): e4318.<br><br>Rooney, B., et al. "SNAPSHOT USA 2019–2023: The first five years of data from a coordinated camera trap survey of the United States." In Press (2024). |
| Carolina Critters | 4260 | Lasky, Monica, et al. "CAROLINA CRITTERS: a collection of camera trap data from wildlife surveys across North Carolina."Â EcologyÂ 102.7 (2021): e03372. |
| Recreational Effects on Wildlife | 1894 | Kays, Roland, et al. "Does hunting or hiking affect wildlife communities in protected areas?."Â Journal of Applied EcologyÂ 54.1 (2017): 242-252. |
| CDFW Bobcat Program | 1368 | Roberts, R. California Department of Fish and Wildlife, Bobcat Program Initiative. wildlifeinsights.org (2023). |
| Urban to Wild Project | 996 | Forrester, T. 2000. Last updated December 2023. Urban to Wild Project. http://n2t.net/ark:/63614/w12004302. Accessed via wildlifeinsights.org on 2024-08-29. |
| NEON | 767 | McMurry et al in prep. NEON camera trapping |
| Okaloosa S.C.I.E.N.C.E. Project | 501 | Forrester, T. 2011. Last updated January 2023. Okaloosa S.C.I.E.N.C.E. Project. http://n2t.net/ark:/63614/w12004287. Accessed via |

| | | |
|---|---|---|
| | | wildlifeinsights.org on 2024-08-29. |
| Albany Area Camera Trapping Project | 155 | Kays, R. 2008. Last updated March 2024. Albany Area Camera Trapping Project. http://n2t.net/ark:/63614/w12003860. Accessed via wildlifeinsights.org on 2024-08-29. |
| ForestGEO | 99 | Jansen, P. (2014) Smithsonian Environmental Research Center ForestGEO Project. http://n2t.net/ark:/63614/w12004305. |
| Smithsonian Grassland Ecology Program | 92 | McShea, W. 1999. Last updated April 2023. Smithsonian Grassland Ecology Program. http://n2t.net/ark:/63614/w12004765. Accessed via wildlifeinsights.org on 2024-08-29. |
| Museums Connect Mexico | 51 | Malleshappa, V., Smithsonian, E., Kays, R., Schuttler, S. (2015). Last updated December 2022. Museums Connect Mexico. http://n2t.net/ark:/63614/w12004298. Accessed via wildlifeinsights.org on 2024-08-29. |
| Calloway Forest Preserve | 49 | McMurry, S., Parsons, A., Lasky, M., Luongo, K., Clark, J., McShea, W., Scher, L., Kays, R., Spurlin, J., Martin, G., Frech, G., Barajas-Salazar, K., Snider, M. (2022). Last updated October 2023. Calloway Forest Preserve. http://n2t.net/ark:/63614/w12004251. Accessed via wildlifeinsights.org on 2024-08-29. |
| Pilot Mountain | 31 | Kays, R., Snider, M., McMurry, S., Alyetama, M. (2024). Last updated April 2024. Pilot Mountain Density 2024. http://n2t.net/ark:/63614/w12007160. Accessed via wildlifeinsights.org on 2024-08-29. |
| Thinking outside the park | 29 | Herrera, Daniel J., et al. "Thinking outside the park: Recommendations for camera trapping mammal communities in the urban matrix."Â Journal of Urban EcologyÂ 7.1 (2021): juaa036. |
| Shingle Shanty Adirondack Mammal Survey | 19 | Kays, R. . 2011. Last updated January 2023. Shingle Shanty Adirondack Mammal Survey. http://n2t.net/ark:/63614/w12004780. Accessed via wildlifeinsights.org on 2024-08-29. |
| Smithsonian - Shenandoah National Park | 18 | Rooney, Brigit; McShea, William. 2023. Last updated August 2024. Shenandoah National Park. http://n2t.net/ark:/63614/w12006220. Accessed via wildlifeinsights.org on 2024-08-29. |
| Collapse of invasive competitor | 15 | McDonald BW, Lashley MA, Cove MV. (2024). Collapse of invasive competitor expands distribution |

| | | |
|---|---|---|
| | | of endangered ecosystem engineer. Global Ecology and Conservation. |

### Table S2: List of modeled species

| Common name | Scientific name | Number of camera trap deployments in modeled range | Number of 10-day CT replicates per deployment (median, 10th quantile - 90th quantile) | Total number of 10-day CT replicates containing an observation of the species | Number of iNaturalist detections of the species | Lineage included? |
|---|---|---|---|---|---|---|
| American badger | *Taxidea taxus* | 5027 | 5 (3-7) | 242 | 1868 | |
| American black bear | *Ursus americanus* | 12764 | 3 (2-7) | 4103 | 23907 | Yes |
| Black-tailed jackrabbit | *Lepus californicus* | 3102 | 5 (3-7) | 1505 | 10160 | |
| Bobcat | *Lynx rufus* | 15880 | 3 (2-7) | 3774 | 16210 | Yes |
| California ground squirrel | *Otospermophilus beecheyi* | 1622 | 5 (4-6) | 573 | 24491 | Yes |
| Collared peccary | *Pecari tajacu* | 387 | 6 (3-8) | 271 | 6470 | |
| Coyote | *Canis latrans* | 16089 | 3 (2-7) | 10652 | 43223 | |
| Desert cottontail | *Sylvilagus audubonii* | 2033 | 5 (3-7) | 646 | 19373 | |
| Douglas's squirrel | *Tamiasciurus douglasii* | 1265 | 5 (4-7) | 1033 | 5754 | Yes |
| Eastern chipmunk | *Tamias striatus* | 1779 | 5 (3-7) | 952 | 37179 | Yes |
| Eastern cottontail | *Sylvilagus floridanus* | 12702 | 3 (2-6) | 3942 | 56738 | |
| Eastern gray squirrel | *Sciurus carolinensis* | 5249 | 3 (2-5) | 7991 | 114932 | Yes |
| Elk | *Cervus canadensis* | 1248 | 5 (3-7) | 651 | 19812 | |
| Fisher | *Pekania pennanti* | 2181 | 4 (3-7) | 273 | 1617 | |

| | | | | | | |
|---|---|---|---|---|---|---|
| Fox squirrel | *Sciurus niger* | 11466 | 3 (2-6) | 3227 | 58385 | |
| Golden mantled ground squirrel | *Callospermophilus lateralis* | 1456 | 5 (3-7) | 323 | 8792 | |
| Grey fox | *Urocyon cinereoargenteus* | 14942 | 3 (2-6) | 3164 | 13341 | Yes |
| Grey wolf | *Canis lupus* | 794 | 6 (3-7) | 120 | 1097 | |
| Groundhog | *Marmota monax* | 9902 | 3 (2-6) | 459 | 18455 | |
| Moose | *Alces alces* | 961 | 5 (3-7) | 175 | 14812 | |
| Mountain cottontail | *Sylvilagus nuttallii* | 1552 | 5 (3-7) | 283 | 1278 | |
| Mule deer | *Odocoileus hemionus* | 3436 | 5 (3-7) | 4802 | 69659 | Yes |
| Nine-banded armadillo | *Dasypus novemcinctus* | 2724 | 4 (2-7) | 2140 | 14331 | |
| North american porcupine | *Erethizon dorsatum* | 3869 | 5 (3-7) | 216 | 10673 | |
| Northern flying squirrel | *Glaucomys sabrinus* | 3022 | 5 (3-7) | 243 | 579 | |
| Northern raccoon | *Procyon lotor* | 15764 | 3 (2-7) | 15580 | 56432 | |
| Pronghorn | *Antilocapra americana* | 891 | 4 (2-6) | 374 | 9160 | |
| Puma | *Puma concolor* | 2829 | 5 (4-7) | 452 | 3770 | |
| Red fox | *Vulpes vulpes* | 10468 | 3 (2-6) | 2784 | 22754 | Yes |
| Red squirrel | *Tamiasciurus hudsonicus* | 7392 | 3 (2-7) | 1467 | 33192 | Yes |
| Snowshoe hare | *Lepus americanus* | 4167 | 5 (3-7) | 577 | 6512 | Yes |
| Southern flying squirrel | *Glaucomys volans* | 12392 | 3 (2-6) | 219 | 1770 | Yes |
| Striped skunk | *Mephitis mephitis* | 14688 | 3 (2-6) | 1683 | 14444 | Yes |
| Virginia opossum | *Didelphis virginiana* | 13411 | 3 (2-6) | 7086 | 23565 | |
| Western gray squirrel | *Sciurus griseus* | 1057 | 5 (4-7) | 952 | 7696 | |
| White-tailed antelope squirrel | *Ammospermophilus leucurus* | 600 | 5 (4-7) | 149 | 3129 | Yes |

| | | | | | |
|---|---|---|---|---|---|
| White-tailed deer | *Odocoileus virginianus* | 14420 | 3 (2-7) | 35871 | 128838 |

### Table S3: Simulation parameters

Parameter values used during simulation for the validation exercise. Note that $\beta'_i$ and $\beta''_i$ are simulated as fixed values rather than random draws from a normal distribution to ensure that realized grouping effects are comparably different from one another across simulations.

| **Parameter** | **Interpretation** | **Simulation value** |
|---|---|---|
| $\beta'_i$ | Group-level SVCs of lineage | Sequence from -1.5 to 1.5 |
| $\beta''_i$ | Group-level SVCs of ecoregion | Sequence from -1.5 to 1.5 |
| $\sigma^2_{CAR}$ | Variance of each CAR-type SVC | 1 |
| $\sigma^2_{\epsilon}$ | Variance of the additive CAR spatial field on intensity | 0.5 |
| $\alpha_{w0}$ | Cloglog-scale detection intercept | -1.5 |
| $\beta_0$ | Log-scale intensity intercept | -1.2 |
| $\boldsymbol{\alpha}_w$, $\boldsymbol{\alpha}_x$, $\beta_{i,0}$ | The spatially stationary components of the window-level effects on detection, the spatial covariate effects on detection, and the spatial covariate effects on intensity | Random draws from unif(-1, 1) |
| $\sigma_m$ | The standard deviation of the location-level random effect on detection | 0.2 |
| $\phi$ | The overdispersion parameter for the iNaturalist negative binomial | 0.1 |
| $\theta_0$ | The log-scale average detection rate in iNaturalist per unit effort | -5 |

| | | |
|---|---|---|
| $\theta_1$ | The slope between iNaturalist per-unit detection rate and animal intensity | 0.5 |
